# Direct inference of the distribution of fitness effects of spontaneous mutations from recombinant inbred *C. elegans* mutation accumulation lines

**DOI:** 10.1101/2024.05.08.593038

**Authors:** Timothy A. Crombie, Moein Rajaei, Ayush S. Saxena, Lindsay M. Johnson, Sayran Saber, Robyn E. Tanny, José Miguel Ponciano, Erik C. Andersen, Juannan Zhou, Charles F. Baer

**Affiliations:** Department of Biology, University of Florida, Gainesville, FL 32611, USA; Department of Molecular Biosciences, Northwestern University, Evanston, IL 60208, USA; Florida Institute of Technology, Melbourne, FL 32901, USA; Department of Biostatistics, Yale School of Public Health, New Haven, CT 06510; Dyne Therapeutics, Waltham, MA 02451, USA; Present address: Florida International University, Miami, FL; Department of Biology, Johns Hopkins University, Baltimore, MD 21218, USA; University of Florida Genetics Institute, Gainesville, FL, 32611, USA

## Abstract

The distribution of fitness effects (DFE) of new mutations plays a central role in evolutionary biology. Estimates of the DFE from experimental Mutation Accumulation (MA) lines are compromised by the complete linkage disequilibrium (LD) between mutations in different lines. To reduce LD, we constructed two sets of recombinant inbred lines from a cross of two *C. elegans* MA lines. One set of lines (“RIAILs”) was intercrossed for ten generations prior to ten generations of selfing; the second set of lines (“RILs”) omitted the intercrossing. Residual LD in the RIAILs is much less than in the RILs, which affects the inferred DFE when the sets of lines are analyzed separately. The best-fit model estimated from all lines (RIAILs + RILs) infers a large fraction of mutations with positive effects (∼40%); models that constrain mutations to have negative effects fit much worse. The conclusion is the same using only the RILs. For the RIAILs, however, models that constrain mutations to have negative effects fit nearly as well as models that allow positive effects. When mutations in high LD are pooled into haplotypes, the inferred DFE becomes increasingly negative-skewed and leptokurtic. We conclude that the conventional wisdom - most mutations have effects near zero, a handful of mutations have effects that are substantially negative and mutations with positive effects are very rare – is likely correct, and that unless it can be shown otherwise, estimates of the DFE that infer a substantial fraction of mutations with positive effects are likely confounded by LD.

## INTRODUCTION

The distribution of fitness effects (DFE) of new mutations is of fundamental importance in numerous areas of evolutionary biology (Fisher 1930; Orr 2000; Peck *et al*. 1997; Schultz and Lynch 1997; Zhang *et al*. 2004), as well as having practical applications, including human genetic disease (Agarwal *et al*. 2023; Boyle *et al*. 2017; Eyre-Walker 2010; Morrow and Connallon 2013) and cancer (Cannataro *et al*. 2016; Cannataro and Townsend 2018; Durrett *et al*. 2010). The DFE can be estimated from data in two ways: indirectly from patterns of sequence variation within and between species (Boyko *et al*. 2008; Gilbert *et al*. 2021; James *et al*. 2023; Johri *et al*. 2020; Keightley and Eyre-Walker 2010; Kim *et al*. 2017; Kousathanas and Keightley 2013; Loewe and Charlesworth 2006; Tataru *et al*. 2017), or directly from comparisons between genotypes differing by a known (or estimated) set of mutations (Böndel *et al*. 2019; Davies *et al*. 1999; Keightley 1994; Ramani *et al*. 2012; Shen *et al*. 2022; Thatcher *et al*. 1998). Each method has strengths and limitations.

Estimation from the standing variation incorporates a vastly larger number of mutations than could ever be assessed experimentally, the effects of very weak selection are detectable (at least in aggregate), and effects are integrated over the entire spectrum of environmental and genomic contexts experienced by the organism in question. However, the method has several important limitations. First, the effects of selection must be jointly estimated with the effects of demography, which are necessarily greatly simplified for analytical tractability (Johri *et al*. 2020; Keightley and Eyre-Walker 2007; Li *et al*. 2012). Second, there is little information about the tail of the distribution for which selection is strong on an evolutionary timescale but weak over the course of a few generations (*s* ≈ 1%) (Kousathanas and Keightley 2013). Third, the method assumes there is a class of mutations that are selectively neutral to serve as a reference; the extent to which that assumption is met is an empirical issue requiring independent validation (Kruglyak *et al*. 2023; Shen *et al*. 2022). Finally, there is no way to connect the DFE back to phenotypic traits.

Direct estimation from fitness differences between known genotypes has the advantage of being conceptually unambiguous - if two groups differ by a single mutation and differ in fitness by some amount *y*, the effect of the mutation is *y*. Constructing two populations that differ by one or a few mutations is straightforward: known mutations can be introgressed or otherwise engineered (e.g., by CRISPR) into a common genetic background to provide “nearly isogenic lines” (NILs). Recent advances in CRISPR technology have made it possible to engineer large panels of NILs in yeast and other microbes (Sharon *et al*. 2018; Shen *et al*. 2022). However, constructing enough NILs to provide a meaningful estimate of the DFE remains a daunting proposition in multicellular organisms. Single-gene “knockout panels”, in which genes are systematically inactivated and the fitness effects documented, have been tremendously important in informing our understanding of the functional aspects of the genome (e.g., Kim *et al*. 2010; Ramani *et al*. 2012; Thatcher *et al*. 1998), but knockout mutations constitute only a small part of the mutational spectrum and do not provide an unbiased estimate of the DFE.

Mutation accumulation (MA) experiments, in which spontaneous mutations are allowed to accumulate in the (near) absence of natural selection, provide the opportunity to estimate the DFE of a (nearly) unbiased set of mutations (Halligan and Keightley 2009; Katju and Bergthorsson 2019). However, within an MA line, all mutations are in complete linkage disequilibrium, which renders individual mutational effects inestimable.

Here we employ a classical line-cross strategy with MA lines, to break down the linkage disequilibrium among the accumulated mutations. We then combine whole-genome sequencing with high-throughput competitive fitness assays to estimate the DFE of a set of 169 spontaneous mutations. This strategy was first employed by Böndel *et al*. (2019) with the unicellular green alga *Chlamydomonas reinhardtii*. We crossed two parental *C. elegans* mutation accumulation (MA) lines derived from the same genetically homogeneous ancestor to get F1 hybrids that are segregating at all mutant loci. The F1s were reciprocally crossed, and from the F2s we constructed two sets of recombinant inbred lines (**Supplemental Figure 1**). For the first set, F2s were further crossed prior to inbreeding to construct a set of Recombinant Inbred Advanced Intercross Lines (RIAILs). For the second set, we omitted the intercrossing step and proceeded directly to the inbreeding step; these lines are classical RILs. We refer to the full set of lines as RI(AI)Ls for brevity. RI(AI)Ls were assayed for competitive fitness against a marked competitor strain nearly isogenic for the ancestral genome, and multilocus genotypes inferred by whole-genome sequencing at low (2-3X) coverage. The strategy is conceptually analogous to QTL analysis, except the variant loci are not simply markers, but rather are the QTL themselves.

## METHODS

### 1. Experimental Methods

#### 1.1 Mutation Accumulation (MA) lines

The details of the MA experiment have been reported elsewhere (Baer *et al*. 2005). Briefly, 100 replicate lines were initiated from a single, highly inbred N2 strain hermaphrodite, and propagated under standard laboratory conditions for a maximum of 250 generations by transfer of a single immature hermaphrodite at four-day intervals. Under this protocol the effective population size, *N_e_* ≈ 1, and all but the most highly deleterious mutations are effectively neutral. The progenitor (G0) was cryopreserved at the outset of the experiment, and surviving MA lines were cryopreserved upon culmination of the MA phase.

#### 1.2 Recombinant Inbred (Advanced Intercross) Lines

Two MA lines (MA530, n=76 mutations and MA563, n=93 mutations) were chosen as parents for a set of recombinant inbred advance intercross lines (RIAILs) or simple recombinant inbred lines (RILs). The parental lines were chosen on the basis of their near-average decline in lifetime reproductive success (∼20%) over four assays after 200 and 220 generations of MA at two different assay temperatures (20° and 25° (Baer *et al*. 2006). The original plan was to construct a set of 600 RIAILs, with ten generations of intercrossing followed by ten generations of selfing, using the “random pair mating with equal contributions of each parent” design of Rockman and Kruglyak (2008; see their Figure 1). However, many crosses failed during the intercrossing phase, so we abandoned the intercrossing and completed the set of lines with RILs. The final set of 517 genotyped lines includes 192 RIAILs and 325 RILs. Details of the crossing schemes are given in **Section I of the Supplemental Material**.

**Figure 1.**
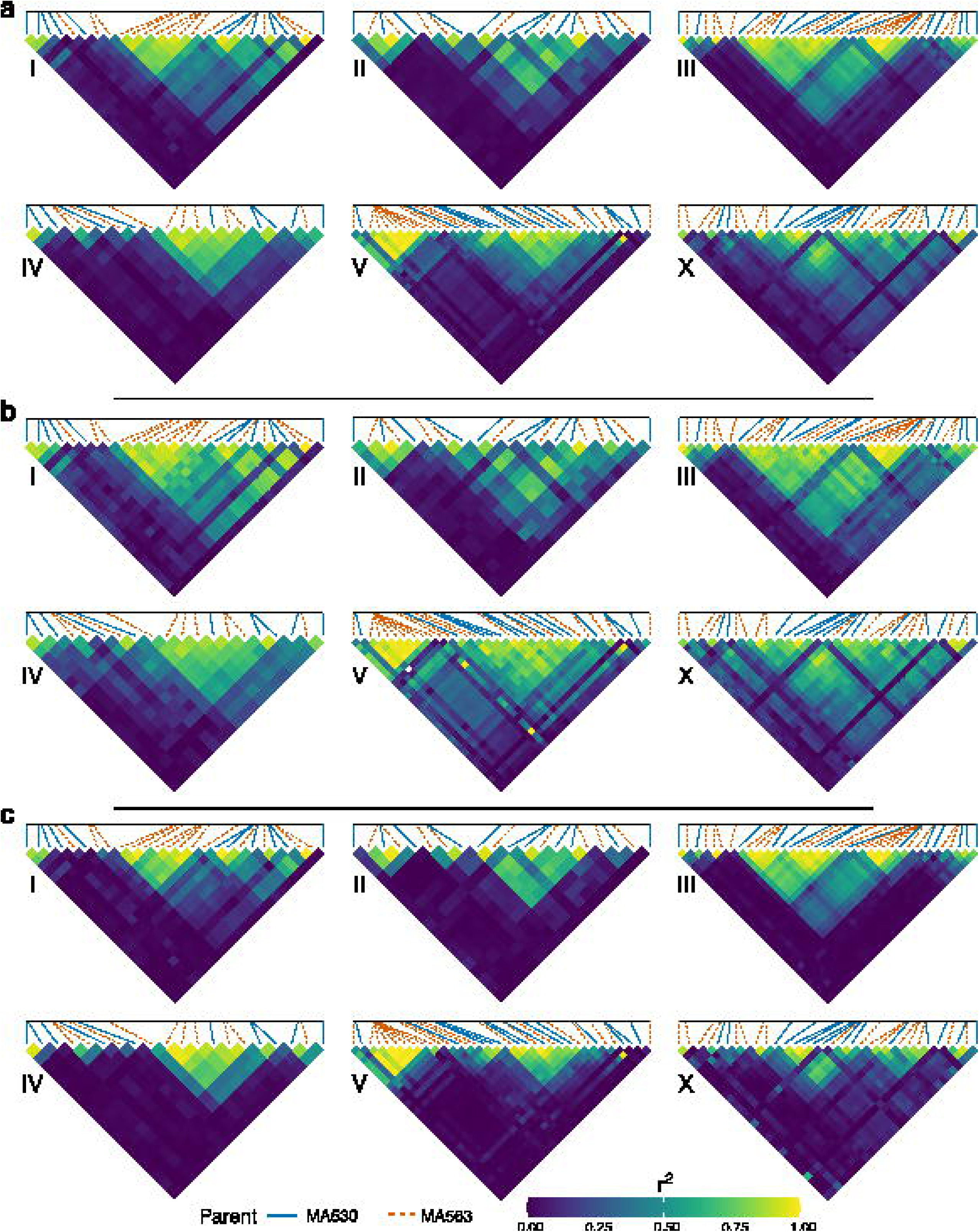
- Intrachromosomal pairwise linkage disequilibrium (LD) (**a**) Pairwise LD (*r^2^*) calculated with all lines (RIAILs + RILs), (**b**) RILs only, and (**c**) RIAILs only. Each heat map represents a chromosome with pairwise LD (*r^2^*) between mutant loci colored as shown in the legend. The colored lines above each chromosome represent the parental origin of the mutant allele (MA530-solid blue, MA563-dashed orange). These lines also show the relative physical position of mutant loci across each chromosome; the far-left vertical line represents the first mutant locus on the chromosome and the far-right vertical line represents the last mutant locus.

#### 1.3 Competitive fitness assays

To assay competitive fitness, an L1-stage focal strain worm and an L1 GFP-marked competitor (strain VP604) were placed together on a plate seeded with bacterial food and allowed to reproduce. Upon exhaustion of the bacterial food, worms were washed from the plate and counted using a Union Biometrica BioSorter™. The natural logarithm of the ratio of the frequencies of the two types, *W*=log[(*p*/1-*p*)], is proportional to the difference in fitness between the focal strain (frequency = *p*) and the competitor strain (frequency = 1-*p*) (Latter and Sved 1994). The assay is described in detail in Appendix 1 of Yeh *et al*. (2018) and summarized in **Section II of the Supplemental Material**.

#### 1.4 Genome sequencing, variant calling, and genotyping

RI(AI)L genomes were sequenced at low (∼2-3X) coverage with 150-bp paired-end Illumina sequencing, using standard methods. Details of sequencing and variant calling are given in **Section III of the Supplemental Material**. Raw sequence data (fastq) of the RI(AI)Ls have been deposited in the NCBI SRA under project number PRJNA1083210. Genome sequences for the G0 progenitor and the parent MA lines have been previously reported (Rajaei *et al*. 2021; Saxena *et al*. 2019).

#### 1.5. Imputation

Given the low (2-3X) sequencing coverage, approximately 1/3 of the data (35.2%) are missing, i.e., the genotype at a given locus was not called as either homozygote. The mean number of loci successfully genotyped per RI(AI)L is 109, and the mean number of RI(AI)Ls for which a locus was scored is 335. To account for the missing genotype information, we constructed a computational procedure to impute the missing data by leveraging linkage disequilibrium (LD; see next section) between segregating sites. Specifically, we used the masked language modeling approach from natural language processing to build a predictive model for the missing alleles. The imputation model is built on the transformer architecture, which has been widely used for modeling natural languages as well as biological sequences such as DNAs and proteins (Ji *et al*. 2021; Rives *et al*. 2021). The model output consists of the predicted log-probability for all possible states per site, i.e., the MA530 or MA563 allele. The details of the model are given in **Section IV of the Supplemental Material**.

To assess the model’s performance, we performed 100 rounds of validation. For each round, all RI(AI)L genotypes were used for training, but with one percent of the called alleles randomly masked. Across the 100 rounds, we observed a high imputation accuracy on the masked positions: mean ± 1 SD prediction accuracy = 90.3 ± 1.5%. Cases in which the imputed allele differs from the called allele include errors in the initial call, so 90% is a conservative estimate of the true prediction accuracy. The final imputed genotypes (**Supplemental Table 1**) were generated by retraining the model on all RI(AI)L genotypes using all available allele information.

#### 1.6. Linkage Disequilibrium (LD)

Alleles from the two parents, MA530 and MA563, are initially in complete coupling (positive) linkage disequilibrium in the F1. However, mutant alleles occur in both parental genomes, so although the initial LD between pairs of mutant alleles is complete, the sign of the association (positive or negative) depends on which parental genomes the mutations occurred. Measures of LD that do not account for the sign of the association are agnostic with respect to whether alleles are coded by the parent of origin or as ancestral (0) vs. mutant (1); the value is the same either way. Measures of LD that do account for the sign of the association may differ by sign depending on if the alleles are coded by parent of origin vs. ancestral vs. mutant. For our purposes, it is more meaningful to code alleles as ancestral or mutant.

The pairwise coefficient of linkage disequilibrium, *D*=*p_A1B1_-p_A1_p_B1_*where *p_A1B1_* is the frequency of the double-mutant (*A1B1*) haplotype at the A and B loci, *p_A1_* is the frequency of the mutant allele at the A locus and *p_B1_* is the frequency of the mutant allele at the B locus. The expected allele frequency in the RI(AI)Ls is 0.5 at all segregating loci, but the observed frequencies will vary due to sampling. We report two measures of LD, the squared coefficient of correlation, *r^2^*, and *D\**=*D/|Dmax*|, where |*Dmax*|= min[*p_A1_*(1-*p_B1_*)], (1-*p_A1_*)*p_B1_*]; *r^2^* is constrained non-negative and *D\** can take on values [-1,1]. Note that our *D\** is the familiar *D’* but with the sign retained. We calculated *r*^2^ and *D\** among all pairs of the 169 loci using the PLINK v1.9 commands ‘--r2’ and ‘--r dprime-signed’ respectively (Purcell *et al*. 2007). We also report the mean pairwise intra-chromosomal and inter-chromosomal LD for (1) all lines (n = 517), (2) RILs only (n = 325), and (3) RIAILs only (n = 192). To visualize intra-chromosomal pairwise LD we used the ggplot2 package v3.4.4 for R Statistical Software v4.2.3 (Wickham 2009).

#### 1.7. Heritability

We estimated the broad-sense heritability (H^2^) of *W* from the among-line (i.e., among-RI(AI)L) component of variance estimated from the general linear model (GLM) *y_ijk_* = μ *+* α*_i_ +* β*_ij_ +* ε*_ijk_*, where *y_ijk_*is the value of *W*, μ is the overall mean, α*_i_*is the random effect of Block *i*, β*_ij_* is the random effect of Line *j* in Block *i*, and ε*_ijk_* is the residual effect of Replicate *k* of Line *j* in Block *i*. Because the RI(AI)Ls are homozygous lines derived from a cross of homozygous parents, V_G_ = V_L_, where V_L_ is the among-line component of variance (Falconer 1989, Ch. 15) and the broad-sense heritability H^2^=V_G_/V_P_, where V_P_ is the total phenotypic variance. Variance components were estimated by restricted maximum likelihood (REML), as implemented in the MIXED procedure of SAS v. 9.4. 95% Confidence intervals of H^2^ were determined empirically from 200 bootstrap replicates, resampling lines pooled over blocks while retaining the effect of Block in the analysis.

To account for the possibility that some of the among-line variance was due to factors other than genotype, we included a set of six “pseudolines” of the G0 ancestor and of each parental MA line in each assay block, which are the experimental equivalent of RILs except they are genetically homogeneous, and any among-(pseudo)line variance must be due to causes other than variation among genes. Pseudolines were analyzed identically to the RI(AI)Ls.

We next estimated the proportion of the total broad-sense heritability <u>not</u> explained by the cumulative additive effects of the mutations, H^2^* (here “additive” formally means “homozygous non-epistatic”, because we have no information about dominance). First, we calculated the multiple regression *y_ijk_* = *μ* **+ *β*x** + *ε*, where *y_ijk_* is the value of *W* as before, *μ* is the overall mean, **x** is the vector of genotypes at mutant loci 1-169, *β* is the vector of regression coefficients, and *ε* is the residual effect. We then re-estimated the linear model from above, *y*_ijk_* = *μ* + *α_i_* + *β_ij_* + *ε_ijk_*, where the terms are as before, where the *y*_ijk_* are the residuals of the multiple regression of *W* on the multilocus genotype, **x**. The difference H^2^-H^2^*\** is the narrow-sense heritability *h^2^*, i.e., the fraction of the total phenotypic variance explained by the additive effects of the mutations. Statistical significance of *h^2^*was assessed by randomly permuting estimates of *W* among replicates and re-calculating *h^2^*.

### 2. Estimation of the DFE

#### 2.1. Raw Difference

The simplest way to measure the phenotypic effect of a mutation at locus *i* is from the average difference in the trait between lines that have the mutant allele and lines that have the ancestral allele at locus *i*. Following Böndel *et al*. (2019) we refer to the mutational effects calculated in this way as the raw difference, *u_RAW_*. Confidence intervals and approximate standard errors of *u_RAW_* were calculated from 1000 bootstrap replicates, holding the number of lines in each category (mutant, wild-type) constant in each (re)sample.

#### 2.2. Bayesian MCMC

We take a fully Bayesian approach to estimate the posterior distribution of all genetic and non-genetic parameters. The basic model is the same as in section 1.7 above, such that the observed fitness of replicate *k* of line *j* in block *i* is: *y_ijk_* = *μ + α_i_ + β**^T^*****x_j_** *+* ε*_ijk_*. The vector *β* contains the effects for the 169 mutations. We fit a series of models with increasing complexity in the prior distribution of *β*, to test different hypotheses regarding the DFE of the mutations. In all models, the grand mean, μ, follows an uninformative normal distribution with mean zero and SD = 10. The individual block effects follow normal distributions with mean 0 and SD = 1, given the small variation in block effects when averaged over lines (SD = 0.13). The models tested are summarized in **Table 1**.

**Table 1.**
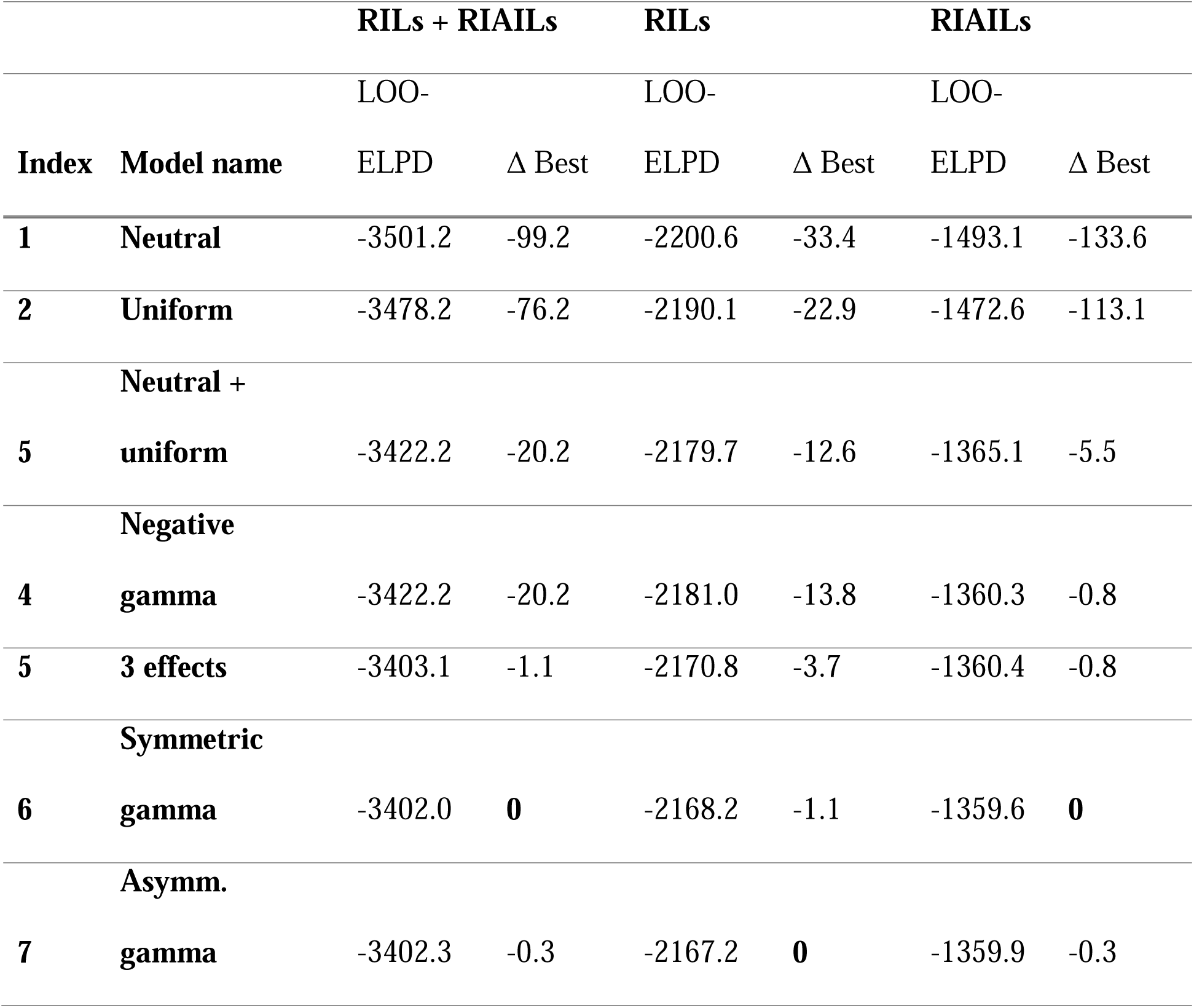
Comparison of seven competing Bayesian models fitted to the genotype and phenotype data of all RILs and RIAILs, and separately to RILs and RIAILs. Each model was run with 50 random genotype replicates. Each replicate consisted of four Markov Chains with 4000 Metropolis steps. Sampling was performed using the software PyMC3 (Salvatier *et al*. 2016Salvatier *et al*. 2016). Model performance is measured using the Bayesian leave-one-out expected log pointwise predictive density (LOO-ELPD), quantifying the generalizability of the fitted model to validation data points. Higher (less negative) LOO-ELPD indicates better model performance.

To begin, in model 1 (“neutral model”) mutational effects are constrained to 0, i.e., *β* = 0. In model 2 (“uniform effect model”), all mutations in the vector *β* have a constant effect (*u*), such that *y_ijk_* = *μ + α_i_ + m_j_ × u + ε_ijk_*., where *m_j_*is the number of mutant alleles in line *j*. Model 3 (“neutral + uniform effect model”) assumes that mutations in vector β follow identical independent distributions such that the *m*-th mutation, *β_m_*, has a probability 1 – *q* of being neutral, and *q* of having a nonzero constant effect *u*, such that *β_m_ = w × u*, where *w* is sampled from a Bernoulli distribution with parameter *q*, which in turn is drawn from an uninformative Beta prior with shape parameter = 2. In both the uniform effect model and the neutral + uniform effect model, the constant mutational effect *u* follows a normal prior with mean 0 and SD = 10. Model 4 (neutral + uniform positive effect + uniform negative effect, “3-effect model”) in addition assumes that mutations can take both constant positive or negative effects, such that *β_m_ = w × (z × u^+^ – (1 – z) × u^-^)*. Similarly, *w* is a Bernoulli random variable with the probability *q*, equal to the probability that a mutation is non-neutral, which follows the same distribution as model 3. The parameter *z* controls the conditional probability of a nonneutral mutation having the positive effect, and is a Bernoulli random variable with probability *p^+^*, which follows an uninformative Beta distribution with shape parameters = 2. The constant positive/negative effects *u_pos_*_/_*u_neg_* follow an uninformative normal distribution with mean 0 and SD = 10.

In addition to these constant-effects models, we tested three models in which mutational effects are sampled from a continuous Gamma distribution. In model 5 (“negative gamma”), all mutations are assumed to have negative (i.e., deleterious) effects, with effect sizes sampled identically and independently from a Gamma distribution, whose shape and rate parameters follow uninformative half normal distributions (SD = 10). In model 6 (“symmetric gamma”) and model 7 (“asymmetric gamma”), mutations can have either positive or negative effects, such that we can express individual mutation effects as β_m_ = *z* × β_m_ ^+^ – (1 – *z*) × β_m_^−^. Similar to model 4, *z* is a Bernoulli random variable with probability *p^+^*, which follows a symmetric Beta distribution. The positive (negative) effect sizes, β_m_ ^+^ (β_m_^−^) are in turn sampled from their respective Gamma distributions, as in Model 5. The only difference between model 6 and 7 is that in model 6, β_m_^+^ and β_m_^−^ follow the same Gamma distribution, whereas in model 7, the Gamma distributions for the positive and negative effect sizes are allowed to be different.

Bayesian inference for all models was implemented in the statistical software PyMC3 v5.10 (Salvatier *et al*. 2016). The No-U-Turn-Sampler was employed to acquire posterior samples. Continuous random variables were sampled using the Hamiltonian Monte Carlo method which relies on gradients calculated using automatic differentiation, whereas discrete random variable were sampled using the Metropolis algorithm. To account for the uncertainty in the genotypes due to missing alleles, for each model we performed 50 independent Monte Carlo runs, each with missing alleles sampled from independent Bernoulli distributions with probability predicted by the trained imputation model. For each model and genotype replicate, we ran 4 parallel Monte Carlo chains, each with 1000 warm up steps and 4000 sampling steps. We used the R-hat statistic (Vehtari *et al*. 2021) as a diagnostic of model divergence, which compares the parameter estimates between and within chains. R-hat is greater than 1 if the chains are not well mixed, such that the between and within-chain sample distributions disagree.

We used a Bayesian model selection procedure to identify the best model. Specifically, for each model we estimated the leave-one-out expected log pointwise predictive density (ELPD LOO) model fit, equal to the mean expected log likelihood of the observed fitness of a random individual given its genotype, calculated based on a model fitted using the full data set minus the focal individual. The procedure is implemented in PyMC3 based on the approximate method introduced by Vehtari *et al*. (2017) The ELPD LOO scores for all 50 genotype replicates were averaged to provide an overall goodness-of-fit score for each model.

## RESULTS

### Linkage Disequilibrium

The purpose of constructing RI(AI)Ls is to break up the linkage disequilibrium between mutations, to permit estimation of the effects of individual mutations. That effort was only partially, and variably, successful. Averaged over all lines (RILs + RIAILs), intrachromosomal LD as measured by median *r*^2^ is 0.12 (**Figure 1; Supplemental Figure 2**). However, LD is much higher in the RILs (median *r*^2^ = 0.28) than in the RIAILs (median *r*^2^ = 0.045). Ten generations of advanced intercrossing was effective in breaking up LD, on average, but regions of near-complete LD remain even in the RIAILs. Inspection of **Figure 1** reveals that regions of high LD are concentrated in the chromosome centers, as expected given the reduced rate of crossing over in centers relative to arms, although there are also regions of high LD in chromosome arms where mutations are tightly clustered. Interchromosomal LD is near 0 in both RILs and RIAILs (**Supplemental Figure 3**), indicating a trivial role for sampling variance in maintaining LD.

### Heritability

Our goal is to estimate the effects of spontaneous mutations on fitness. To begin, we ask: is there heritable variation in competitive fitness among the RI(AI)Ls? The broad-sense heritability of *W* including all RI(AI)Ls, H^2^=0.30 (bootstrap 95% CI=0.271, 0.370). Estimates of H^2^ were similar for RIAILs (H^2^=0.337; bootstrap 95% CI=0.256, 0.403) and RILs (H^2^=0.313; bootstrap 95% CI=0.243, 0.382). Including all RI(AI)Ls, narrow-sense heritability, estimated from the residuals of the multiple regression of *W* on multilocus genotype, *h^2^* = 0.16 (permutation test, P<0.001; averaged over 1000 permutations of the data, random *h^2^* = 0.023, max=0.048). The cumulative additive effects of the 169 segregating spontaneous mutations explain approximately half of the total heritable variance in *W*. By way of comparison, H^2^ for competitive fitness from a set of 28 *C. elegans* wild isolates was 0.49, although the assays in the two studies are not directly comparable (Teotónio *et al*. 2006).

Considering RIAILs and RILs separately, *h^2^* of the RILs is similar to the estimate from the full dataset (*h^2^* = 0.20, n=325), whereas the same analysis for RIAILs gives a REML point estimate of residual V_L_=0. Taken at face value, these results imply that additive mutational effects completely explain H^2^ (i.e., *h^2^* = H^2^) in the RIAILs, whereas the additive effects only explain about two-thirds of the among-line variance in the RILs. To investigate the possibility that LD could explain the unexplained among-line variance in the RILs, we used parametric bootstrap simulations, as follows. For each RIL we (*i*) assigned each mutation in its genome a fitness effect drawn from a given DFE with mean effect equal to the observed mean, (*ii*) summed the effects across loci, and (*iii*) added to each replicate a residual (= microenvironmental) fitness effect drawn from a normal distribution. We then estimated H^2^ and *h^2^* from the simulated data as described above. In the first set of simulations (n=100), we maintained the observed LD structure; in the second set of simulations we permuted alleles (mutant or ancestral) among loci in each RIL to break up the LD. We tested two different DFEs. The first DFE is the ‘asymmetric Gamma model’ described in Methods, where mutations can have positive or negative effects, with the magnitude of the positive/negative effect drawn from two non-identical Gamma distributions. The second DFE is the ‘negative gamma’ model, where mutations can only have negative effects and are drawn from a single Gamma distribution. We sampled effects of mutations from these two DFEs using the posterior mean model parameters (Supplemental Table 2). Residual fitness effects were sampled from zero-mean normal distributions with variance equal to the posterior means of the noise variance inferred jointly with model parameters for the two DFEs (σ^2^ ≈ 1). For both DFEs, LD had no effect on the inferred *h^2^*; in each case *h^2^*=H^2^ in 100% of the simulations, as expected because the mutations were the only source of among-line variance in the simulations.

Having ruled out differences in LD as the cause of missing heritability in the RILs if mutational effects are strictly additive, the remaining unexplained heritability in the RILs must be due to some combination of epistasis, transgenerational epigenetic inheritance (TEI), and/or residual (but small) genotype-environment correlations. It is not obvious at first glance why the same set of epistatic mutations would lead to missing heritability in the RILs but not in the RIAILs. However, the number of RIAILs (n=192) is only slightly greater than the number of loci (n=169), so it is plausible that there simply is little power to detect residual among-line variance once the additive effects of the mutations are accounted for. When *h^2^* is estimated for the full set of RI(AI)Ls with the additive effects regressed separately for each block, the residual heritability disappears; that result reinforces the likelihood that the absence of missing heritability in the RIAILs is simply due to lack of power rather than an actual absence of non-additive among-line variance. We elaborate on this possibility in Section V of the Supplemental Material.

To account for potential non-genetic variation that is nevertheless heritable over a few generations, we estimated variance components among sets of “pseudolines” of the G0 ancestor of the parental lines, and of the MA530 and MA563 parental lines. These controls are not powerful (n=30 pseudolines, 6 per block), but in all three cases the REML estimate of the among-pseudoline component of variance, V_L_= 0.

### Relationship between number of mutations and mean fitness

If all mutational effects are equal and in the same direction (i.e., the Bateman-Mukai criteria (Mukai 1964)), the slope of the regression of *W* on the number of mutant alleles carried by a line will equal the average effect of a mutation. Averaged over all RI(AI)Ls, accounting for variation among assay blocks and removing two outlying lines, the regression of *W* on number of mutations is not significantly different from 0 (slope = −0.0051, F_1,509_=1.83, P>0.17), although the trend suggests that mutations are deleterious, on average.

### Relationship between mutational effect and mutant allele frequency

The expected frequency of segregating neutral alleles in the RI(AI)Ls is 0.5. Selection was minimally effective in the crossing and inbreeding phases (*N_e_* ≈ 2), but it was not absent. If most mutations are deleterious and if deleterious alleles were preferentially removed by selection, then (*i*) the average frequency of mutant alleles will be < 0.5, and (*ii*), there should be a negative relationship between allele frequency and mutational effect size. The mean observed mutant allele frequency is 0.500 (range = 0.287-0.675). The correlation between mutant allele frequency *p_i_* at the *i*th locus and the raw difference *u_RAW,i_*, *r_pu_*= 0.15 (**Supplemental Figure 4**). Thus, we infer that selection did not systematically skew mutant allele frequencies away from the expected neutral frequency.

### The Bayesian posterior DFE

To infer the DFE, we tested a series of seven increasingly complex models, using the Bayesian MCMC analysis outlined in the Methods. Because of the discrepancy in average LD between the RIAILs and the RILs, all analyses were first done on the full set of RI(AI)Ls, and repeated on RIAILs and RILs separately.

As a first step, we tested for model convergence, using the R-hat statistic. We observed no divergence between the four parallel Markov chains, indicated by R-hat < 1 in all cases (Vehtari *et al*. 2021). Model performance, as measured by the Bayesian leave-one-out expected log pointwise predictive density (LOO-ELPD, Vehtari *et al*. 2021) averaged across 50 genotype replicates, is summarized in **Table 1**. Posterior means and 95% credible intervals of model parameters are given in **Supplemental Table 2**.

#### (i) All lines (RILs + RIAILs = RI(AI)Ls)

Reassuringly, the **neutral** model, in which mutational effects are constrained to equal 0, performs worst. The **uniform** effect model, in which mutational effects are constrained to be equal, is moderately better (Δfit = 23.0). The posterior mean for the shared mutational effect (*u*) is negative and has a 95% credible interval not intersecting zero (*u* = −0.006; CI = −0.009, −0.005).

The **neutral + uniform effect model**, in which mutations can either have a uniform non-zero effect with probability *q* or be neutral with probability *1-q*, performed significantly better (Δfit = 50.0). Again, the mean mutational effect is inferred to be negative (*u* = −0.16, 95% CI = - 0.24, −0.10), but with low probability (*q* = 0.064, 95% CI = 0.026, 0.114). The **negative Gamma** model, in which effects are constrained to be negative and sampled from a Gamma distribution, fits equally well as the neutral + fixed effect model (*u* = −0.007, Δfit = 0.0).

All models summarized so far assume mutations must have a uniform sign. The first model relaxing this assumption is the **3-effect model**, in which a mutation can be neutral with probability 1 – *q,* or have a fixed positive/negative effect with probabilities *q^+^* and *q^-^* (in our Bayesian model parametrization, *q^+^ = q × p^+^, q^-^ = q × (1 – p^+^)*, where *p^+^* is the probability that a mutation has a positive effect, given that it is non-neutral). This model showed a significant improvement in performance (Δfit = 19.1).

Finally, the two-sided Gamma models (**symmetric** and **asymmetric Gamma**) provide a moderate improvement over the 3-effect model. The two models have LOO-ELPD scores that are nearly indistinguishable (symmetric gamma model = −3402, asymmetric gamma model = - 3402.3), indicating that the additional flexibility conferred by the asymmetric gamma model does not confer higher generalizability to new data. For the asymmetric gamma model, the alpha (scale) and beta (rate, inverse of the scale parameter) parameters for the positive and negative halves of the distribution have nearly identical posterior distributions (**Supplemental Table 2**). Additionally, the two-sided gamma models show very similar posterior distributions for all parameters. We therefore focus our discussion on the more parsimonious symmetric gamma model.

On average, mutations are slightly less likely to have a positive effect (*p^+^* = 0.426; 95% CI = 0.294, 0.547). The posterior distribution of the effects of all 169 mutations shows that 39.6% of all mutations have a positive posterior mean effect (**Figure 2A**), consistent with the posterior probabilities *p^+^*/ *^-^*. However, individual mutations exhibit large credible intervals that intersect zero (**Figure 3A**). The distribution of negative mean effects shows a longer tail than the positive effects, but this asymmetry in shape was not reflected in the model selection results, where the symmetric and asymmetric Gamma models have virtually identical performance. This is likely a power issue, whereby the increased flexibility of the asymmetric Gamma model was not supported by enough data to result in likelihood improvements that can offset the penalty resulting from the higher model complexity.

**Figure 2.**
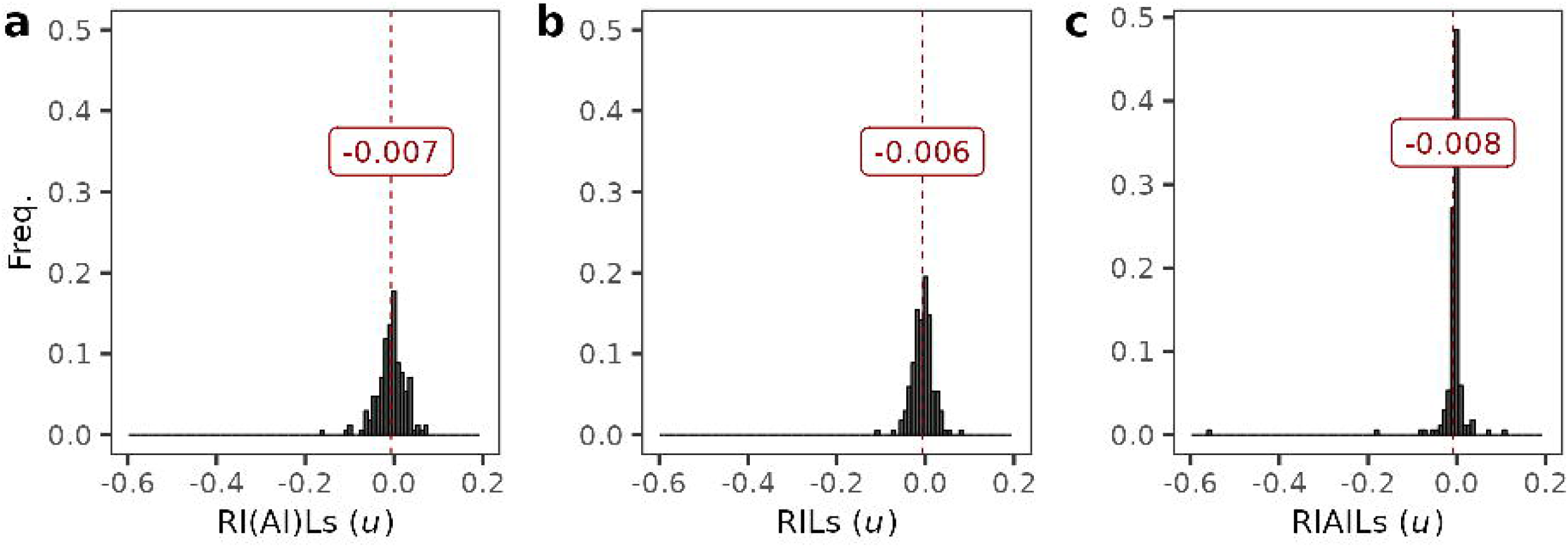
- Distribution of Bayesian posterior mutational effects on fitness. The distribution of mean mutational effects (*u*) calculated using the Bayesian MCMC method is shown. The distribution is calculated separately with (**a**) all lines (**b**), RILs only, or (**c**) RIAILs only. The vertical red line in each panel represents the mean of means for that population. The mean value for each panel is also annotated on the plots in red text.

**Figure 3.**
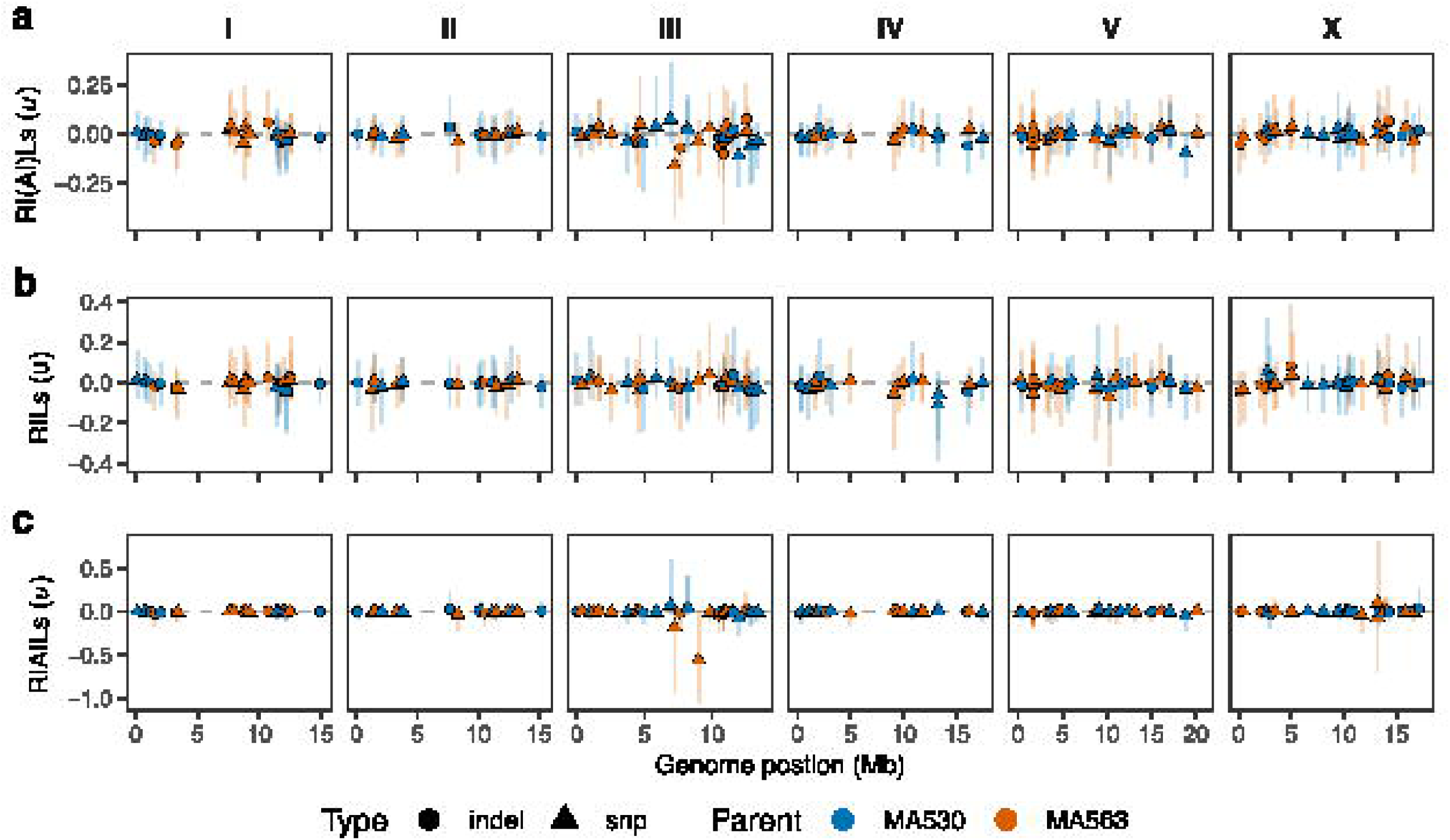
- Bayesian posterior mutational effects by genome position. The mutant loci are plotted by their physical position in the genome (x-axis) and their mean mutational effect (*u*) (y-axis), which was calculated using the Bayesian Markov chain Monte Carlo (MCMC) method. The colors indicate the parent of origin for the mutant locus (MA530-blue, MA563-orange) and the shapes show the mutant class (indel-circle, snp-triangle). The vertical lines plotted behind each point represent the 95% confidence intervals of the mutant effect estimates. The mutational effects are calculated separately with all lines (**a**), RILs only (**b**), and RIAILs only (**c**).

#### (ii) RILs

The model selection results for the RILs are largely consistent with results based on the full set of RI(AI)Ls. The neutral and the fixed effect models have the lowest LOO-ELPD (**Table 1**). The two negative effects models have similar LOO-ELPD values and show significant improvement over the first two models. Finally, we see that the three two-sided models provide further substantial improvement over the one-sided model. The two-sided Gamma models produced very similar LOO-ELPD scores, while the 3-effect model has a moderately lower value. The distribution of mean mutational effects under the symmetric Gamma model are similar to results generated from the full set of RI(AI)Ls (Pearson’s *r* = 0.56; **Figure 2B**).

#### (iii) RIAILs

Model selection results for the RIAILs reveal a different pattern. Although the neutral and fixed effect models still perform worst, performance of the models in which effects are constrained to be non-positive (in particular the negative Gamma model) is now close to that of the two-sided models (**Table 1**). The similarity between the two-sided models and the negative-only model is supported by the change in the shape of the two-sided gamma models, in which the frequency of mutations with positive effects is lower (*q+* = 0.355; 95% CI 0.119, 0.595). Inference from RIAILs resulted in an overall reduction in the mean posterior effects of mutations, such that the effects of most mutations are shrunk towards zero (**Figure 2C**). Additionally, the posterior variance of the mutational effects is lower in the RIAILs (mean posterior SD of mutational effects is 0.040, compared with 0.056 in the full set of RI(AI)Ls) (**Figure 3C**), even with the lower sample size. The mutational effects for the RIAILs are more weakly correlated to those inferred from the full set of RI(AI)Ls (Pearson’s *r* = 0.36) than are the effects inferred from the RILs.

#### (iv) Locus-specific effects

The simplest way to infer the mutational effect at a locus is to calculate the mean value of all lines with a mutant allele and all lines with an ancestral allele at that locus; the difference is the raw difference (**u_RAW_**) of the mutation at that locus. As a sanity check, we plotted the inferred Bayesian posterior effect against the raw difference; ideally, the correlation should be +1. The correlations were positive, but well below 1 in all three cases (**Figure 4**). The magnitude of the raw difference is typically much larger than that of the posterior effects. The difference is likely caused by LD, in that the raw difference of a single mutation contains contributions from other linked mutations, which may inflate the estimates.

**Figure 4.**
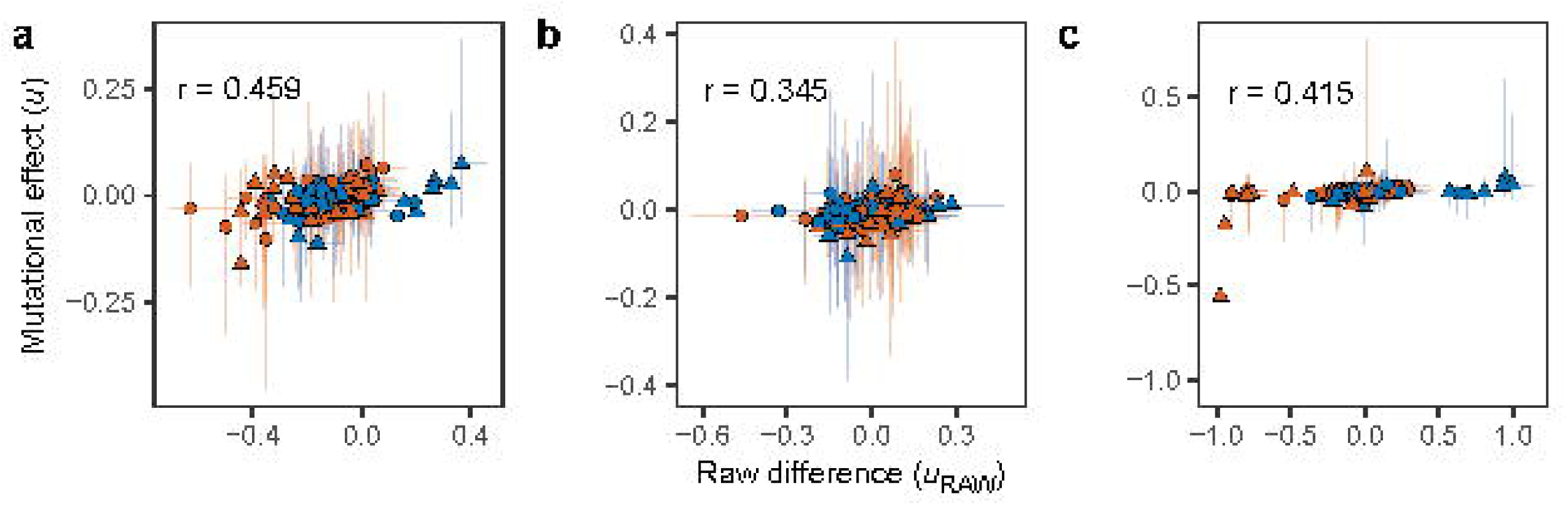
- The relationship between Bayesian posterior mutational effects (*u*) and raw difference, *u_RAW_*. The effects are calculated separately using (**a**) all lines, (**b**) RILs only, or (**c**) RIAILs only. Each point represents a locus and is colored by the parent of origin (MA530-blue, MA563-orange). The shape of the point shows the mutant class (indel=circle, snp=triangle). 95% confidence intervals for the estimates are plotted as vertical and horizontal lines behind the points. Pearson’s correlation coefficient (*r*) is displayed in the upper left of each panel.

### Effects of mutant haplotypes

A major challenge is that many mutations are in high LD, making the effects of individual mutations nearly unidentifiable (for example, if two mutations with effects, *u_1_* and *u_2_* are in complete LD, we only have observations for the sum of their effect *u_1_ + u_2_*, making it impossible to estimate *u_1_* and *u_2_* separately). To proceed, we first identified haplotype blocks consisting of groups of loci in which LD among all pairs of consecutive loci *r*^2^ > 0.8. We then designated two haplotypes for each haplotype block. Among loci in a haplotype block, two types of haplotype assignment can occur. Consider a haplotype block with two loci, each with an ancestral and a mutant allele (coded 0 and 1). If the two loci are in positive LD, we have an ancestral haplotype (00) and a double-mutant haplotype (11). If the two loci are in negative LD, we have two single-mutant haplotypes, 01 and 10. Treating the data as haplotypes rather than individual loci reduces the sample size from 169 (the number of loci) to 114 (the number of haplotypes). We restricted this analysis to the symmetric Gamma model.

We acquired the posterior sample of a mutant haplotype by summing the posterior samples of the individual mutations at each locus in the haplotype. We repeated this procedure for the RILs, RIAILs and the full set of RI(AI)Ls. In all three cases, the distribution of the mean mutant haplotype effects is skewed to the left (**Figure 5**). The percentage of mutant haplotypes with negative posterior means is 61.4% in the full set of RI(AI)Ls, 64.0% in the RILs, and 67.5% in the RIAILs. Again, inference from the RIAILs results in an overall reduction in the mean and variance of posterior effects of mutant haplotypes, relative to inferences from RILs and the full set of RI(AI)Ls. The mean absolute posterior mean effect for the negative mutant haplotypes based on RIAILs only (*u^-^* = −0.022) is twice that of the positive mutant haplotypes (*u^+^* = 0.011).

**Figure 5.**
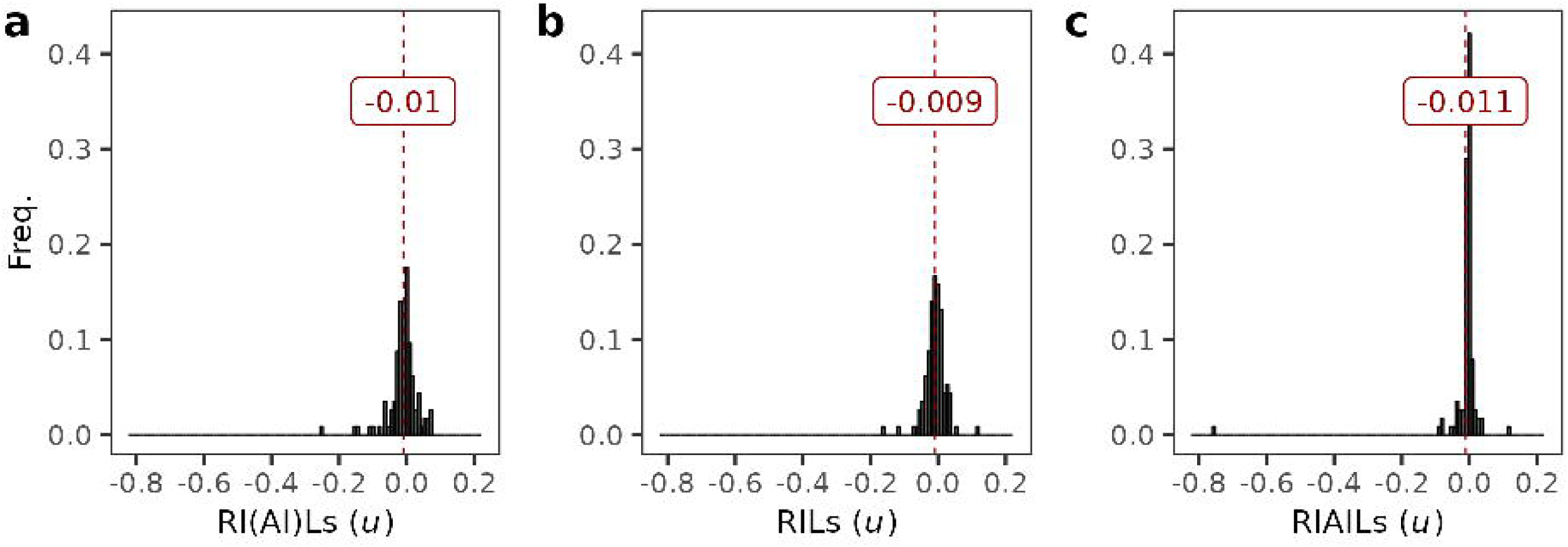
- Distribution of Bayesian posterior mutant haplotype effects on fitness. The distribution of mean mutant haplotype effects (*u*) calculated using the Bayesian MCMC method is shown. The distribution is calculated separately for (**a**) all lines, (**b**) RILs only, or (**c**) RIAILs. The vertical red line in each panel represents the mean of means for that population. The mean value for each panel is also annotated on the plots in red text.

Finally, the lower LD in the RIAILs allowed us to identify a mutant haplotype with a strong negative effect located in a 6.05 Mb region between positions 3771123 and 9819058 on chromosome III (**Figure 6**). This haplotype contains 13 mutations, including 11 SNPs and 2 indels. The two mutant haplotypes are 1000111001100 for MA530, and 0111000110011 for MA563. The MA563 mutant haplotype has a large negative effect (*u* = −0.760; 95% CI −1.09, - 0.149), whereas the MA530 mutant haplotype shows a moderately strong positive mean effect (*u* = 0.118; 95% CI −0.134, 0.647). However, their effects are strongly negatively correlated in the posterior samples, i.e., if an estimated effect at the MA530 haplotype is large and negative, the corresponding estimate at the MA563 haplotype is large and positive. The most we can say with confidence is that the cumulative effect of mutations in this region is to reduce *W* by about 0.64 relative to the ancestor, which is sufficient to explain the decrease in fitness of MA563 relative to the ancestor (**Supplemental Figure 5**).

**Figure 6.**
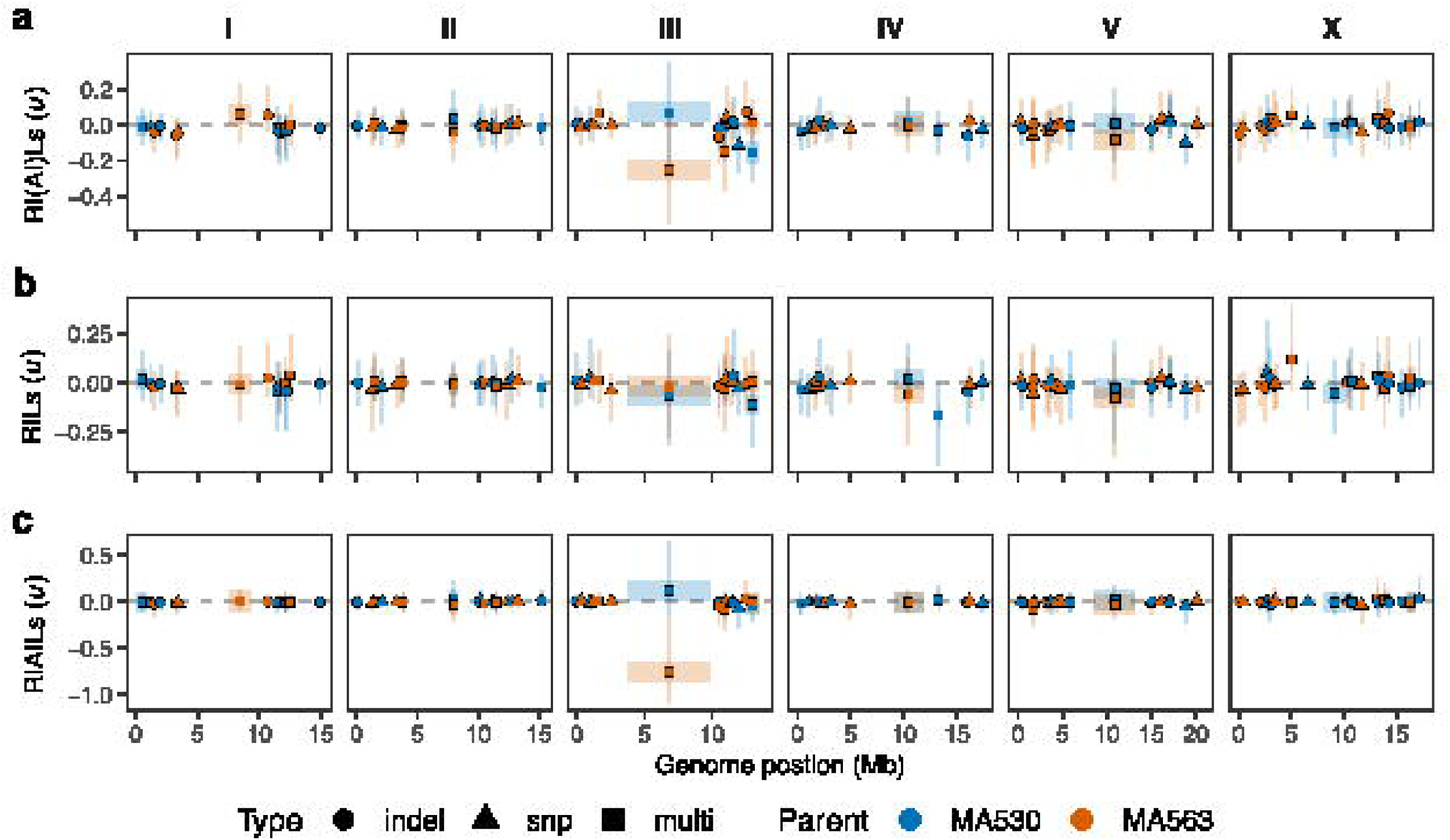
- Bayesian posterior mutant haplotype effects by genome position. The 114 mutant haplotypes are plotted by their physical position in the genome (x-axis) and their mean haplotype effect (*u*) (y-axis), which was calculated using the Bayesian Markov chain Monte Carlo (MCMC) method. The center of haplotypes are plotted as points and the genomic range of multi-locus haplotypes are represented by horizontal boxes plotted behind the points. The colors indicate the parent of origin for the mutant haplotype (MA530-blue, MA563-orange). Multi-locus mutant haplotypes are plotted with square points (multi), and the other single-locus haplotypes are plotted with shapes based on mutation type (indel-circle, snp-triangle). The vertical lines plotted behind each point represent the 95% confidence intervals of the haplotype effect estimates. The haplotype effects are calculated separately with all lines (**a**), RILs only (**b**), and RIAILs only (**c**).

The full list of mutations, along with parent of origin and their inferred effects, are presented in **Supplemental Table 3**; fitness data are presented in **Supplemental Table 4**.

## DISCUSSION

Unsurprisingly, mutations are deleterious, on average. Coincidentally or not, the point estimate of the mean average raw difference in competitive fitness in the RI(AI)Ls, −0.0039, is extremely similar to the same estimate from the full set of 80 MA lines of which the two parental lines were drawn. Assuming that a random pair of MA lines differs by 160 mutations, the average mutational effect estimated from the data of Yeh *et al*. (2018, Table 1) is −0.0040. Given the substantial sources of variation in these experiments, the concordance is remarkable. In a similar vein, Yeh et al. estimated the mutational heritability from the same data, 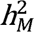 =*V_L_*/2*t* = 0.00084/generation of MA. Summed over the approximately 250 generations of MA, we predict a broad-sense heritability H^2^ ≈ 0.2, about 2/3 of the observed value in this study. Or differently put, our estimate of H^2^ implies a mutational heritability 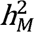 ≈ 0.0012. Given that both measures of heritability are ratios of variances, the observed values are quite consistent.

Perhaps more surprising is the relatively high narrow-sense heritability of the mutational effects (*h^2^*=0.16), which explain roughly half of the heritable variance in fitness. There are no comparable competitive fitness data from wild isolates, but Zhang *et al*. (2021) estimated H^2^ and *h*^2^ for lifetime fecundity on solid media for a set of 121 *C. elegans* wild isolates. In their assay *h*^2^ (0.20) was about 1/3 of H^2^ (0.63). In contrast to our RI(AI)Ls, which differ by about 85 mutations on average, the wild isolates differ by thousands of segregating variants. Comparison of heritabilities is problematic because the upper bound is 1, which means that *h*^2^ necessarily reaches an asymptotic value. However, if we assume that the contribution of non-heritable effects (V_E_) is similar in the two studies – and we would naively expect that V_E_ is greater in a competitive fitness assay than in a non-competitive assay because the competitor contributes to V_E_ – the implication is that the asymptote is reached after at most a few hundred generations of mutations have accumulated in the population.

The inclusion of both RILs and RIAILs in the experiment is fortuitous. If we only had RILs to work with, we would have been much more confident in concluding that a large proportion of mutations have positive effects. Ten generations of intercrossing in the RIAILs broke up most of the initial LD, but not all of it, and it is clear that at least some of the apparently greater fraction of positive-effect mutations in the RILs can be attributed to the confounding effect of negative-effect mutations in LD. Inspection of the DFE along the chromosome (**Figure 3**) reveals a negative spatial autocorrelation: mutations inferred to have large positive effects are usually in close proximity to one or more mutations with large negative effects.

This study was motivated by three antecedents: the studies of Böndel *et al*. (2019), who used a related crossing design to estimate the DFE from spontaneous MA lines in the unicellular green alga *Chlamydomonas reinhardtii*; of Gilbert *et al*. (2021), who estimated the *C. elegans* DFE from the standing site frequency spectrum among wild isolates; and those of Vassilieva *et al*. (2000) and Keightley *et al*. (2000), who estimated the DFE from the distribution of (non-competitive) fitnesses among *C. elegans* MA lines. We consider each in turn.

Böndel *et al*.’s crossing design differed from ours in a key way: they backcrossed MA lines to the common ancestor rather than crossing two MA lines. Their design results in all mutations being initially in complete coupling (positive) LD, rather than a random mix of coupling and repulsion LD, as in our design. Nevertheless, their design is still constrained to infer the cumulative effects of mutations in LD. They did not report LD, nor did they report the distribution of mutational effects along the chromosomes (except as raw data). They too observed a high proportion of mutations with positive effects on fitness; in their best-fit model (two-sided Gamma with different means for positive and negative DFEs), the DFE was highly leptokurtic, with posterior mean frequency of positive effects, *q^+^*, of 84%. However, the estimated mean (absolute) effect of deleterious mutations, *u^-^*, was 4-5 times greater than the mean positive effect, which reconciles the high frequency of mutations with positive effects with the consistent and well-supported overall decline in fitness of the MA lines. They too observed a strong positive correlation between the inferred posterior mean mutational effect at a locus and the raw difference, and that the Bayesian posterior DFE was shrunk toward zero compared to the raw difference.

Gilbert *et al*. used maximum likelihood, as implemented in the DFE-alpha software (Keightley and Eyre-Walker 2007), to infer the DFE from segregating SNP variation in a set of ∼300 *C. elegans* wild isolates. They also analyzed data simulated under realistic parameters of mutation and recombination to investigate the effect of self-fertilization on the inferred DFE. They found that, while DFE-alpha reprises the input DFE quite faithfully when mating is random, self-fertilization biases the results toward mutations of small negative effect, evidently due to the slower decay of LD under selfing. Inclusion of a small fraction (0.1%) of beneficial mutations similarly biases the inferred DFE of deleterious mutations toward small effects.

*C. elegans* MA lines invariably decline in fitness, and early studies concluded that the mean deleterious mutational effect is quite large (∼10-25 %) (Estes *et al*. 2004; Keightley and Caballero 1997; Vassilieva *et al*. 2000), although none of those studies investigated competitive fitness. The point estimate of the mean deleterious mutational effect from our neutral + uniform effect model (Model 3) in the full set of RI(AI)Ls is −0.16 and the inferred fraction of deleterious mutations (0.064) translates to a per-genome, per-generation deleterious mutation rate of U ≈ 0.02, very consistent with the aforementioned studies. Coincidentally or not, our inference from RIAIL haplotypes that the *C. elegans* DFE consists of a very large proportion of mutations with near-zero effects interspersed with a small number of mutations with large negative effects is very similar to the conclusion of Keightley *et al*. (2000), who reached that conclusion from the distribution of fitnesses among *C. elegans* MA lines that had been subjected to EMS mutagenesis.

## Conclusions

– Two primary conclusions emerge from this work. First, mathematics is no substitute for recombination where inference of the DFE is concerned. When mutations are in strong LD – repulsion or coupling – different combinations of positive and negative effects can result in the same cumulative effect, possibly leading to the mistaken inference that the DFE includes a large fraction of mutations with positive effects. However, posterior estimates at linked loci will be strongly negatively correlated, which will not be true of unlinked loci. That conclusion is obvious in hindsight, and should serve as a cautionary note. But second, the unplanned inclusion in this study of RILs along with the RIAILs, and the large difference in average LD between the two sets of lines, turns out to be informative. As LD is reduced in the RIAILs vs. the RILs, the DFE becomes more leptokurtic, the inferred proportion of mutations with negative effects increases, and the relative difference in magnitude between negative and positive effects increases (negative effects become increasingly greater). When mutations are binned into haplotypes, the most intuitive interpretation of the results is that almost all mutations have effects that are very close to 0, and that the decline in fitness with MA is the result of a small number of mutations with large negative effects – perhaps only one, on chromosome III in the MA563 genome.

Looking ahead, we envision understanding of the DFE being advanced in three ways. First, technical advances in high-throughput gene editing will allow efficient construction of nearly-isogenic lines (NILs), removing the confounding effects of LD. The mutation spectrum can be inferred, and a large random sample of spontaneous mutations can be engineered into a common genomic background(s) and the DFE estimated as we have done here. Second, the DFE of a common set of mutations should be estimated in a variety of contexts. We only assayed fitness in one context in this experiment; it would be very interesting to see if, and how, the DFE changes in different contexts. Finally, experimental estimates of the DFE can be employed as strong priors in estimates of the DFE from standing polymorphism, which may have the added benefit of facilitating estimates of demographic parameters by de-confounding selection from demography.

## Supporting information

Supplemental Table 4_fitness

Supplemental Table 1_genotype matrix

Supplemental Table 2_posterior statistics

Supplemental Table 3_list of mutations and effects

## DATA AVAILABILITY STATEMENT

Raw sequence data have been submitted to the NCBI BioProject database (https://www.ncbi.nlm.nih.gov/bioproject/) under accession numbers PRJNA1083210 (RI(AI)Ls) and PRJNA429972 (parental MA lines). Cryopreserved stocks (G0 ancestor, parental MA lines and RI(AI)Ls) are available upon request to CFB. All code for analyses is available at https://github.com/Crombie-Lab/manuscript_DFE/tree/main

## ACKNOWLEGEMENTS

We thank Joanna Dembek and Mike Snyder for expert work in the lab during the line construction and assay phases of the experiment. We thank David McCandlish, Aneil Agrawal, Ian Dworkin, and an anonymous reviewer for comments on the manuscript. Support was provided by NIH awards GM107227 to CFB, ECA, and JMP, and GM127433 to CFB, ECA, and V. Katju.

